# Phytotoxic tryptoquialanines produced *in vivo* by *Penicillium digitatum* are exported in extracellular vesicles

**DOI:** 10.1101/2020.12.03.411132

**Authors:** Jonas Henrique Costa, Jaqueline Moraes Bazioli, Luidy Darllan Barbosa, Pedro Luis Theodoro dos Santos Júnior, Flavia C. G. Reis, Tabata Klimeck, Camila Manoel Crnkovic, Roberto G. S. Berlinck, Alessandra Sussulini, Marcio L. Rodrigues, Taícia Pacheco Fill

## Abstract

*Penicillium digitatum* is the most aggressive pathogen of citrus fruits. Tryptoquialanines are major indole alkaloids produced by *P. digitatum*. It is unknown if tryptoquialanines are involved in the damage of citrus fruits caused by *P. digitatum*. To investigate the pathogenic roles of tryptoquialanines, we initially asked if tryptoquialanines could affect the germination of *Citrus sinensis* seeds. Exposure of the citrus seeds to tryptoquialanine A resulted in a complete inhibition of germination and an altered metabolic response. Since this phytotoxic effect requires the extracellular export of tryptoquialanine A, we investigated the mechanisms of extracellular delivery of this alkaloid in *P. digitatum*. We detected extracellular vesicles (EVs) released by *P. digitatum* both in culture and during infection of citrus fruits. Compositional analysis of EVs produced during infection revealed the presence of a complex cargo, which included tryptoquialanines and the mycotoxin fungisporin. The EVs also presented phytotoxicity activity *in vitro*, and caused damage to the tissues of citrus seeds. Through molecular networking, it was observed that the metabolites present in the *P. digitatum* EVs are produced in all of its possible hosts. Our results reveal a novel phytopathogenic role of *P. digitatum* EVs and tryptoquialanine A, implying that this alkaloid is exported in EVs during plant infection.

**IMPORTANCE:** During the post-harvest period, citrus fruits can be affected by phytopathogens such as *Penicillium digitatum*, which causes the green mold disease and is responsible for up to 90 % of the total citrus losses. Chemical fungicides are widely used to prevent the green mold disease, leading to concerns about environmental and health risks. To develop safer alternatives to control phytopathogens, it is necessary to understand the molecular basis of infection during the host-pathogen interaction. In the *P. digitatum* model, the virulence strategies are poorly known. Here, we describe the production of phytotoxic extracellular vesicles (EVs) by *P. digitatum* during the infection of citrus fruits. We also characterized the secondary metabolites in the cargo of EVs and found in this set of molecules an inhibitor of seed germination. Since EVs and secondary metabolites have been related to virulence mechanisms in other host-pathogen interactions, our data are important for the comprehension of how *P. digitatum* causes damage to its primary hosts.

## INTRODUCTION

Citriculture is a worldwide multi-billion-dollar activity (1). Brazil, China and the United States are the major producers of citrus (2). In Brazil, 230.000 direct and indirect jobs are related to citriculture (3). The citrus industry in Brazil corresponded to US$ 6.5 billion revenues in 2019 (3). Citrus fruits can be affected by different diseases, leading to up to 50 % of fruit losses, and causing a negative economic impact (4–6).

Diseases caused by fungal pathogens are the most adverse factors causing fresh fruit and vegetable losses during the postharvest period (7, 8). Postharvest losses due to fungal diseases can reach up to 30-55 % of production (8–10). The most damaging postharvest disease in citrus is the green mold caused by *Penicillium digitatum*, which accounts for up to 90 % of citrus losses (4–6).

Demethylation inhibitors (DMI) including prochloraz and imazalil, are fungicides used to combat *P. digitatum* (4). However, this practice has raised concerns about its effects on human health, and in the development of antifungal resistance (10, 11). Therefore, developing safer approaches to control postharvest diseases has become a global trend (7, 8, 10, 11). The development of alternative antifungal tools demands an improved knowledge on how *P. digitatum* causes damage to citrus fruits. Most efforts in this direction have focused on the use of biocontrol agents, antagonist microorganisms, and natural products (8, 11, 12) to neutralize virulence factors (4, 13). Nevertheless, the molecular mechanisms underlying the induction of *P. digitatum*-mediated damage in host cells remain poorly known (4, 5, 14).

Secondary metabolites were reported as essential for fungal pathogenicity, and a consequent attenuation of the plant defense responses (4, 5, 14). Siderophores, for instance, have been associated with fungal virulence by iron sequestration (4, 16). Secondary metabolites have not been yet associated to the pathogenic process promoted by *P. digitatum* (4). Tryptoquialanines are major metabolites produced by *P. digitatum* (17). However, their role in *P. digitatum* phytopathogenicity is unclear, even though these compounds displayed insecticidal (18) and antifungal (19) activities.

In addition to secondary metabolites, extracellular vesicles (EVs) have been associated with the pathogenesis of several infectious diseases (20, 21). EVs are spherical structures that are released by bacteria, fungi, and plant cells (20–22). EVs are delimited by a lipid bilayer membrane in association with proteins, lipids, enzymes, pigments, polysaccharides, and RNAs (20–22). In fungi, EVs were first reported in the human pathogenic yeast *Cryptococcus neoformans* and subsequently characterized in several fungal pathogens (21, 22). In host-microbe interactions, EVs are key players determining the pathogenic outcome (23, 24), and mediating the transfer of virulence traits (25). In plant infections, the roles of EVs have been superficially explored, and the knowledge of how EVs impact the plant physiology is limited (23, 24). It is known that under stress conditions, release of EVs by plants is increased in response to infection (23,24). EVs are involved in the defense of plants against pathogens, forming physical barriers or delivering molecules that are toxic to invading microbes (23, 24). On the other hand, pathogen EVs can inhibit plant immune responses through the export of virulence factors (23). So far, no information on the production of EVs by phytopathogens and their association with secondary metabolites cargo is available.

Herein we report the phytotoxic activity of indole alkaloids and EVs produced by *P. digitatum* in germination assays of *Citrus sinensis* seeds. The export of tryptoquialanines in *P. digitatum* involved EVs. Untargeted metabolomics was applied to confirm the EV-mediated metabolite export in the possible hosts (citrus and stone fruits) affected by *P. digitatum*.

## RESULTS

### Phytotoxicity activity of tryptoquialanine

To investigate the pathogenic potential of tryptoquialanine A (TA), we first purified this alkaloid from *P. digitatum*’s crude extracts (Fig. S1). We then evaluated its phytotoxicity in germination assays of *Citrus sinensis* seeds. TA significantly inhibited seed germination at all tested concentrations. Seeds exposed to concentrations of TA lower than 1.000 ppm exhibited a delay in their germination time when compared to the negative control (NC), as evidenced by the changes in seed colors and size (Fig. 1). Seeds treated with the highest concentration of TA (3.000 ppm) showed a stronger phytotoxic effect, and no formation of radicle was observed (Fig. 1).

**FIG 1.**
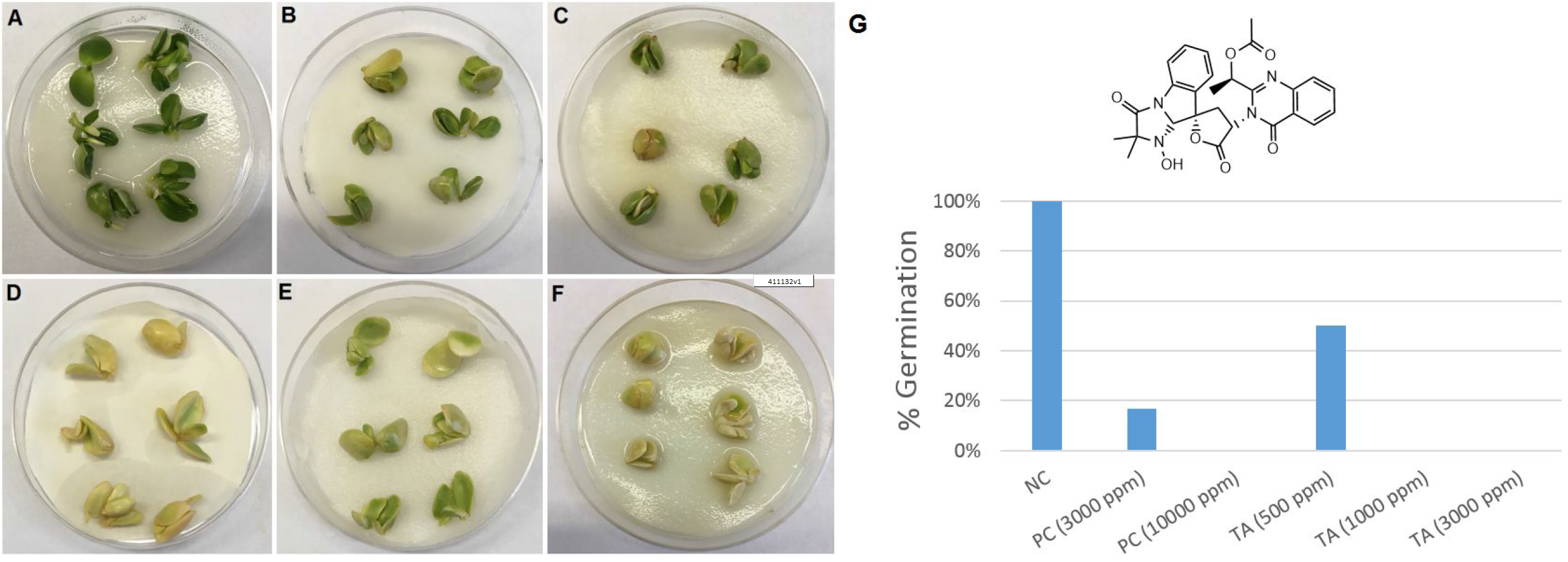
Tryptoquialanine A (TA) is an efficient inhibitor of germination in *C. sinensis* seeds. Untreated seeds (negative control) showed a regular pattern of germination (A). Seeds exposed to TA (500, 1.000, or 3.000 ppm; B, C and D, respectively), or to the commercial herbicide RoundUP® (3.000 or 10.000 ppm; E and F, respectively) manifested defective germination. Seeds exposed to TA (3.000 ppm) and positive control (PC) (10.000 ppm) did not germinate. This visual perception was confirmed by the quantitative determination of germination (%) of *C. sinensis* seeds under different treatments (G). Six seeds were used in each treatment.

Following the observation of the phytotoxic activity of TA against the *C. sinensis* seeds, we compared the metabolite profiles of the seeds exposed to the different treatments by Ultrahigh-pressure liquid chromatography-mass spectrometry (UHPLC-MS). Principal component analysis (PCA) of Quality Control (QC), Negative Control (NC), Herbicide (PC), and Tryptoquialanine A (TA) extracts was performed to observe data reproducibility and grouping tendencies (Fig. 2A). Data reproducibility was verified as QC samples formed a distinct cluster. The two principal components, PC1 and PC2, were responsible for 37.9 % of the variance of the data, revealing a separation between the seed groups related to the treatment received (water, glyphosate herbicide, or TA).

**FIG 2.**
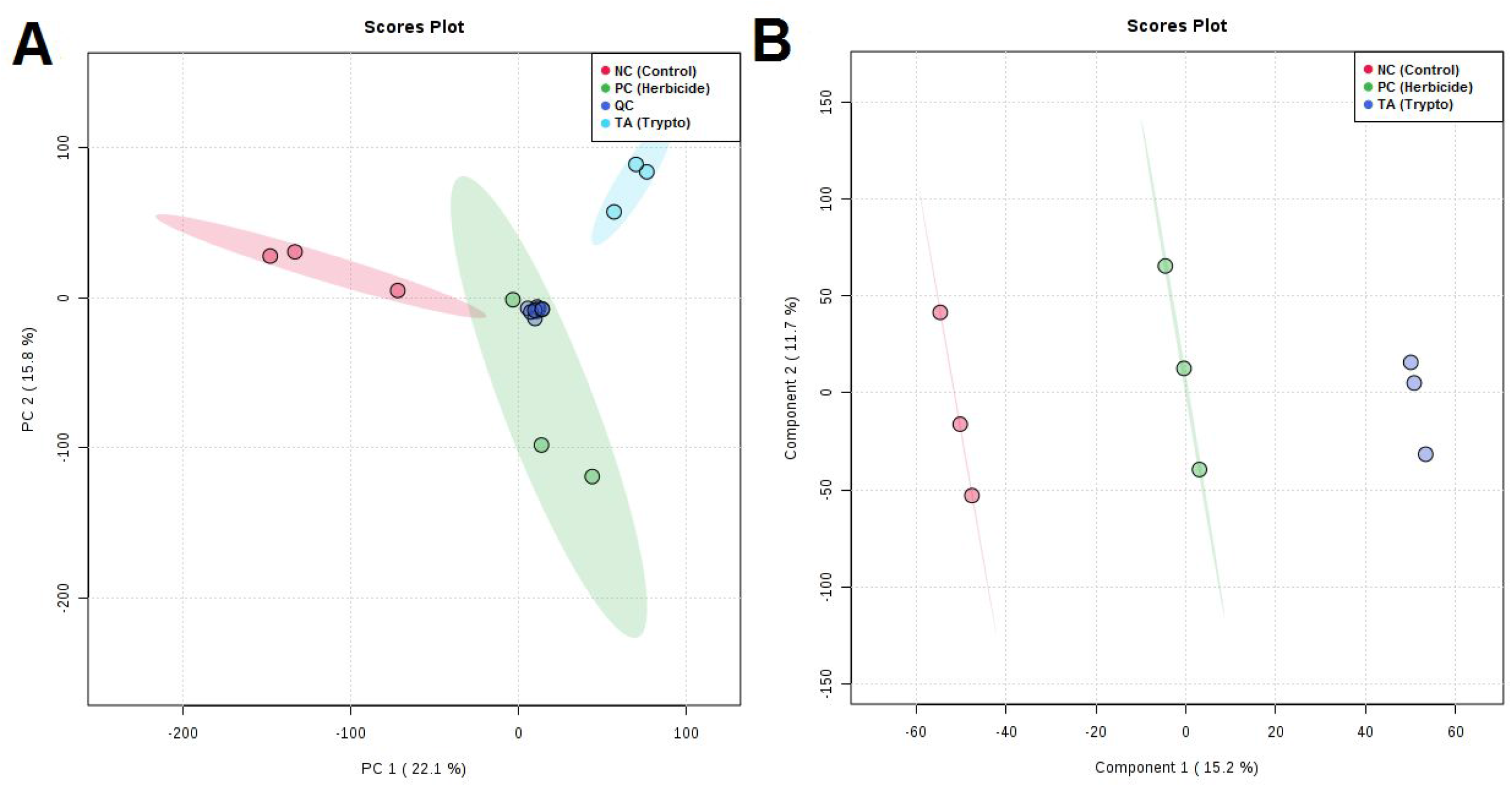
PCA (A) and PLS-DA (B) for extracts of *C. sinensis* seeds treated with water (Negative Control), glyphosate (Herbicide), or tryptoquialanine A (TA).

In order to verify the classification of the seeds according to the treatment received, partial least squares discriminant analysis (PLS-DA) was performed. PLS-DA scores plot confirmed a clear separation between the seed groups (Fig. 2B). In PLS-DA, PC1 and PC2 accounted for 26.9 % of the variance (15.2 % for PC1 and 11.7 % for PC2). As in the PCA scores plot, control seeds were distributed in the opposite way of seeds treated with TA along PC1, while seeds treated with herbicide were plotted in the center.

### *P. digitatum* produces EVs *in vitro* and *in vivo*

To inhibit the germination of *C. senensis* seeds, TA is required to reach the extracellular environment. We then asked if the extracellular export of TA could be vesicle-mediated. However, the production of EVs by *P. digitatum* has not been reported so far. To address this question, we used methods for EV detection in different models of *P. digitatum* growth. Specifically, the production of EVs was evaluated in both solid agar medium and infected citrus fruits. Transmission electron microscopy (TEM) of *P. digitatum* samples grown *in vitro* revealed membranous structures with the typical features of vesicles, including round-shaped structures with bilayered membranes in the 100 nm size range (Fig. 3 A-D). Similar results were observed for vesicles isolated from infected fruits. These results were confirmed by a second experimental approach. Nanoparticle tracking analysis (NTA) of the same samples revealed particles mostly concentrated in the 100-200 nm range, with subpopulations in the 200-300 and 300-400 nm size ranges (Fig. 3 E-F). *In vitro* and *in vivo* samples had similar properties, which were consistent with those previously described for fungal EVs (20, 26, 27).

**FIG 3.**
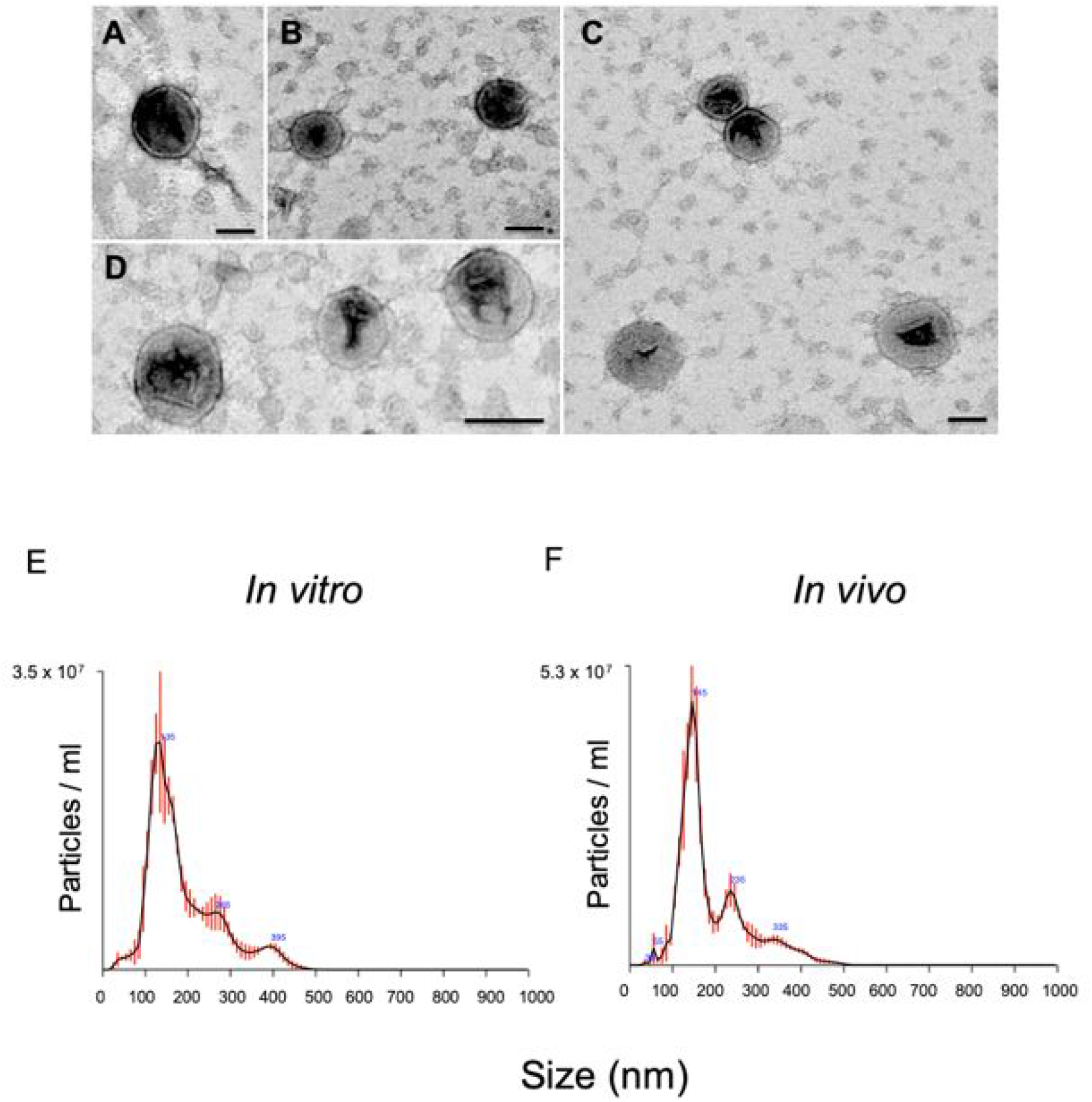
Production of EVs by *P. digitatum*. Analysis of ultracentrifugation pellets by TEM (A-D) revealed the presence of the typical round-shaped structures presenting double membranes. Similar results were obtained with samples obtained *in vivo* (A and B) and *in vitro* (C and D). The visual observations by TEM were confirmed by NTA, which detected particles mostly concentrated in the 100-200 nm range, with subpopulations in the 200-300 and 300-400 nm size ranges. Similar results were obtained with *in vitro* (E) and *in vivo* (F) samples. One representative experiment of three independent replicates producing similar results is illustrated.

### Tryptoquialanine A is a component of *P. digitatum* EVs

The metabolite composition of the *P. digitatum* EVs was investigated by UHPLC-MS/MS in EV extracts obtained *in vivo* (Fig. S2 and S3), followed by molecular networking in the Global Natural Products Social Molecular Networking (GNPS) platform and, when available, compared with standard metabolites. Molecular networking revealed three clusters (A, B, and C) that exhibited compounds present in the EVs (Fig. 4; pink symbols). Metabolites were manually identified by accurate mass analysis, MS/MS fragmentation profiles, comparison with authentic standards (tryptoquialanine A and B) or identified as a hit in the GNPS database. The observed signals corresponded, respectively, to tryptoquialanine A (*m*/*z* 519.19), tryptoquialanine B (*m*/*z* 505.17), deoxytryptoquialanine (*m*/*z* 503.19), *cyclo*-(Phe-Val-Val-Tyr) (*m*/*z* 509.27), Phe-Val-Val-Phe (*m*/*z* 511.29), Phe-Val-Val-Tyr (*m*/*z* 527.28) and *cyclo*-(Phe-Phe-Val-Val) (*m*/*z* 493.28).

**FIG 4.**
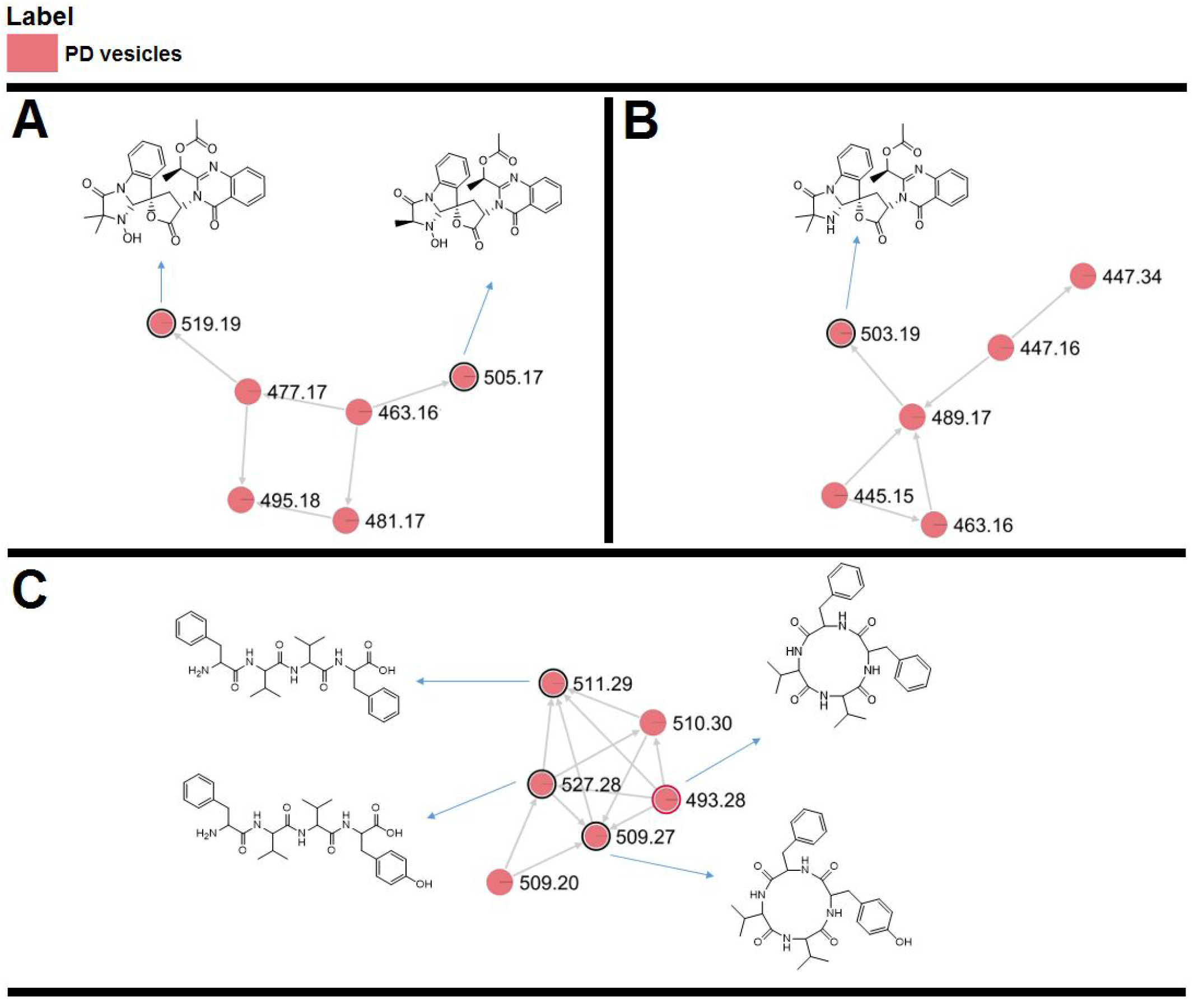
Molecular networking obtained for the *P. digitatum* EV cargo. Indole alkaloids were grouped in clusters A and B, and tetrapeptides were grouped in cluster C. Nodes circled in black indicate molecules identified manually through exact masses and MS/MS fragmentation patterns comparison to the literature. Nodes circled in red indicate molecules identified by comparison with the GNPS platform database.

In the molecular networking analysis, each consensus MS/MS spectrum is represented by a node, and all nodes are labeled with their precursor mass. Indole alkaloids produced by *P. digitatum* were grouped in clusters A and B since they showed similar fragmentation patterns, with typical indole alkaloid fragments observed at [M + H]^+^ *m*/*z* 156.07, *m*/*z* 197.10 and *m*/*z* 213.10 (Fig. S4). Tryptoquialanines A and B and deoxytryptoquialanine are the final products of the tryptoquialanine biosynthetic pathway (28) and, as already mentioned in this section, these indole alkaloids were reported as major secondary metabolites for *P. digitatum* (17). In cluster C, GNPS database indicated the presence of *cyclo*-(Phe-Phe-Val-Val) (Fig. S5A), a mycotoxin known as fungisporin. We also observed that fungisporin analogues were grouped in this cluster. A fragmentation pattern with typical ions observed at [M + H]^+^ *m*/*z* 120.08, *m*/*z* 219.15 and *m*/*z* 247.14 was previously described for compounds Phe-Val-Val-Phe and Phe-Val-Val-Tyr (19, 29, 30) (Fig. S5B and S5C).

### Quantification of tryptoquialanine A in *P. digitatum* EVs

The quantitative composition of alkaloids in *P. digitatum* EVs was evaluated using UHPLC-MS/MS analyses. Firstly, a calibration curve was prepared using standard TA (t_R_ = 7.2 min) (Fig. S1) purified from *P. digitatum*’s crude extracts (18). The coefficient of determination (r^2^) obtained was greater than 0.998, indicating an excellent linearity (Fig. S6). Extracts of *P. digitatum* EVs isolated from *in vivo* assays were again analyzed for the presence of TA. Each 1.0 × 10^10^ *P. digitatum* EVs contained 0.0184±0.0002 µg of TA.

### *P. digitatum* EVs are phytotoxic to seeds

We asked whether the phytotoxic effects of TA alone would be comparable to its vesicle-exported form. To address this question, we isolated EVs produced during infection and performed the seed germination tests in the presence of the vesicles. EVs were adjusted to a final concentration of 2.1 × 10^10^ EVs mL^-1^ to allow comparisons between the effects of purified TA and the vesicle preparations.

After 10 days of incubation, seeds exposed to EVs had germination rates similar to those observed in untreated systems. Positive controls of inhibition of germination revealed seeds with different color and patterns and absence of radicle formation, as expected. However, the seeds that were exposed to the *P. digitatum* EVs showed altered tissues. Tissular alteration included injured areas with differences in pigmentation (Fig. 5). No color alteration or tissue damage were observed in the negative controls.

**FIG 5.**
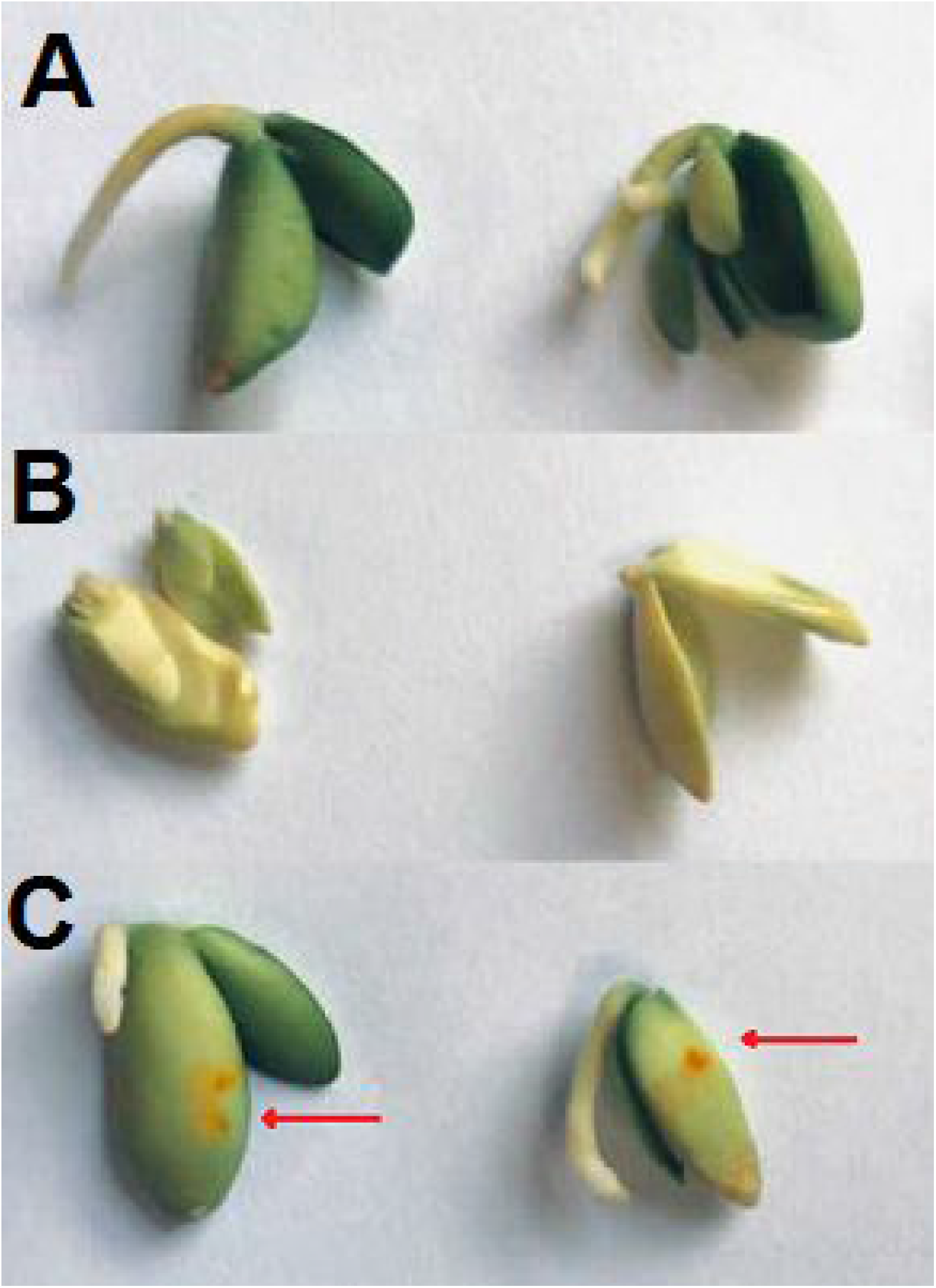
*P. digitatum* EVs affect *C. sinensis* seeds. A) Negative control (NC), consisting of *C. sinensis* seeds incubated in PBS. B) Positive control (PC), consisting of the citrus seeds incubated in glyphosate (10.000 ppm). C) Incubation of *C. sinensis* seeds with *P. digitatum* EVs produced *in vivo* (2.1 × 10^10^ EVs mL^-1^). Seeds exposed to fungal vesicles presented injured tissues (orange spots on their surface; red arrows).

### Comparison of secondary metabolite production of *P. digitatum* in different hosts

After evaluating the citrus response to TA and EVs, we next analyzed the metabolite response of different hosts to the infection caused by *P. digitatum*. Similarly, to what we observed for the *P. digitatum* EV extracts, molecular networking of extracts from plums and oranges infected with *P. digitatum* showed three clusters (D, E and F) with compounds present only in the infected fruits (blue, green and yellow nodes) and absent in control fruits (orange and pink nodes) (Fig. 6 and 7). Metabolites were manually identified by their accurate masses and fragmentation profiles, or identified as hits in the GNPS database. Fragmentation patterns obtained by MS/MS analyses are represented in Fig. S7. Accurate mass measurements showed mass errors below 5 ppm (Table 1). The structures of the detected metabolites are shown in Fig. 8.

**FIG 6.**
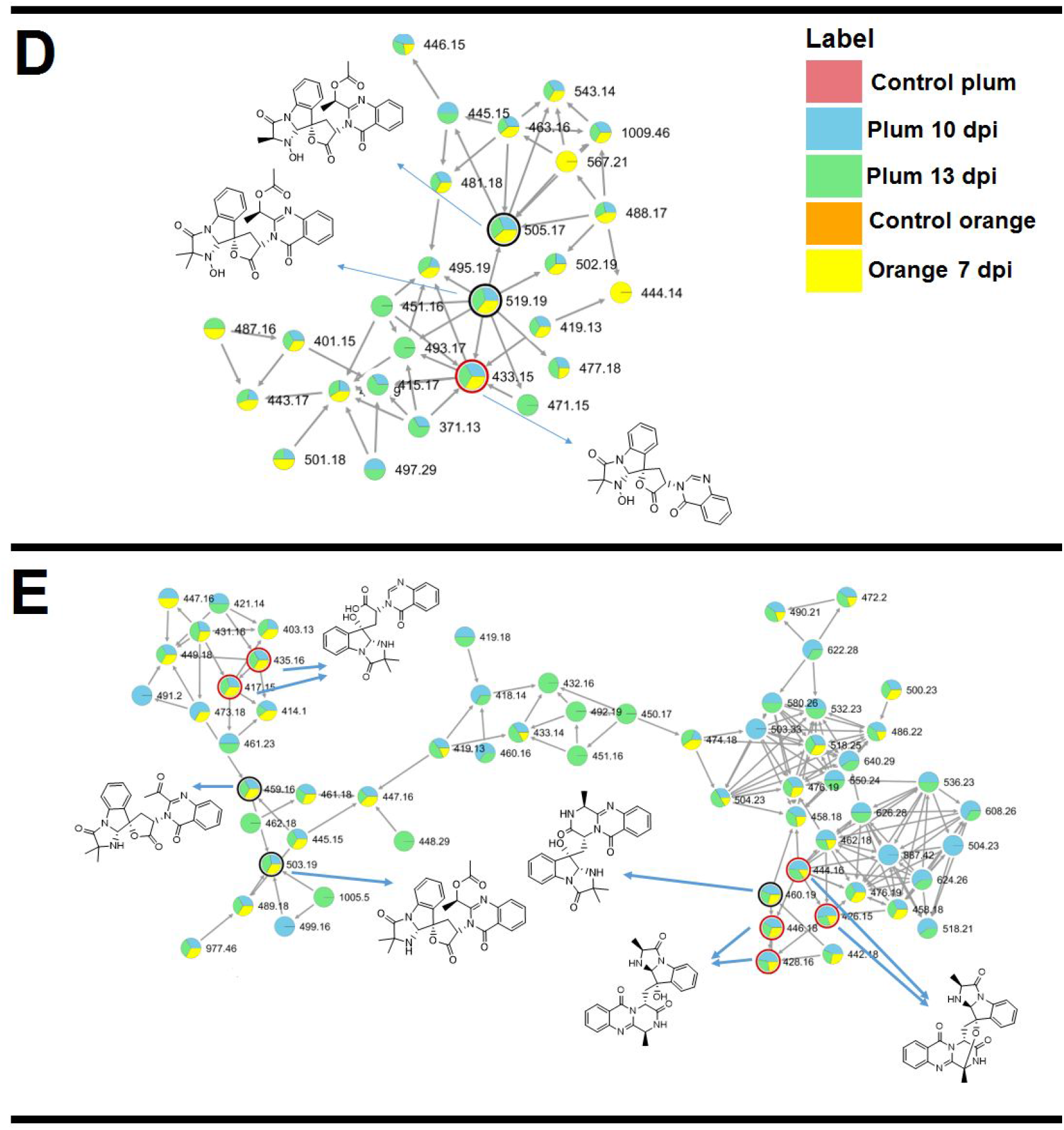
MS/MS molecular network of different extracts of *P. digitatum*. Nodes circled in black indicate molecules identified manually through accurate mass and fragmentation pattern analysis. Nodes circled in red indicate molecules identified by comparison with the GNPS database. Indole alkaloids produced by *P. digitatum* were grouped in clusters D and E.

**FIG 7.**
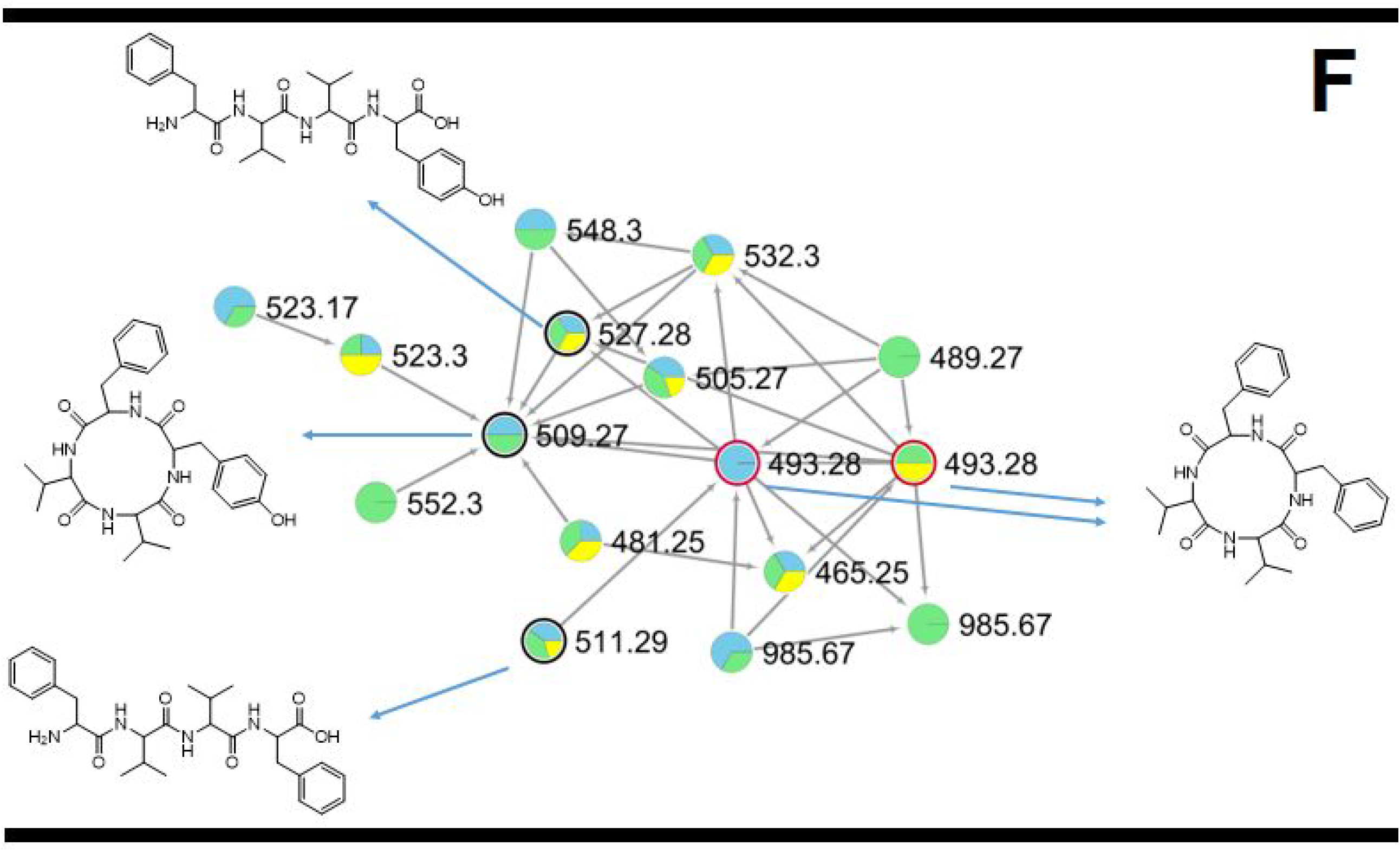
MS/MS molecular network of different extracts of *P. digitatum*. Nodes circled in black indicate molecules identified manually through accurate mass and fragmentation pattern analysis. Nodes circled in red indicate molecules identified by comparison with the GNPS database. Fungisporin and analogues were grouped in cluster F.

**Table 1.**
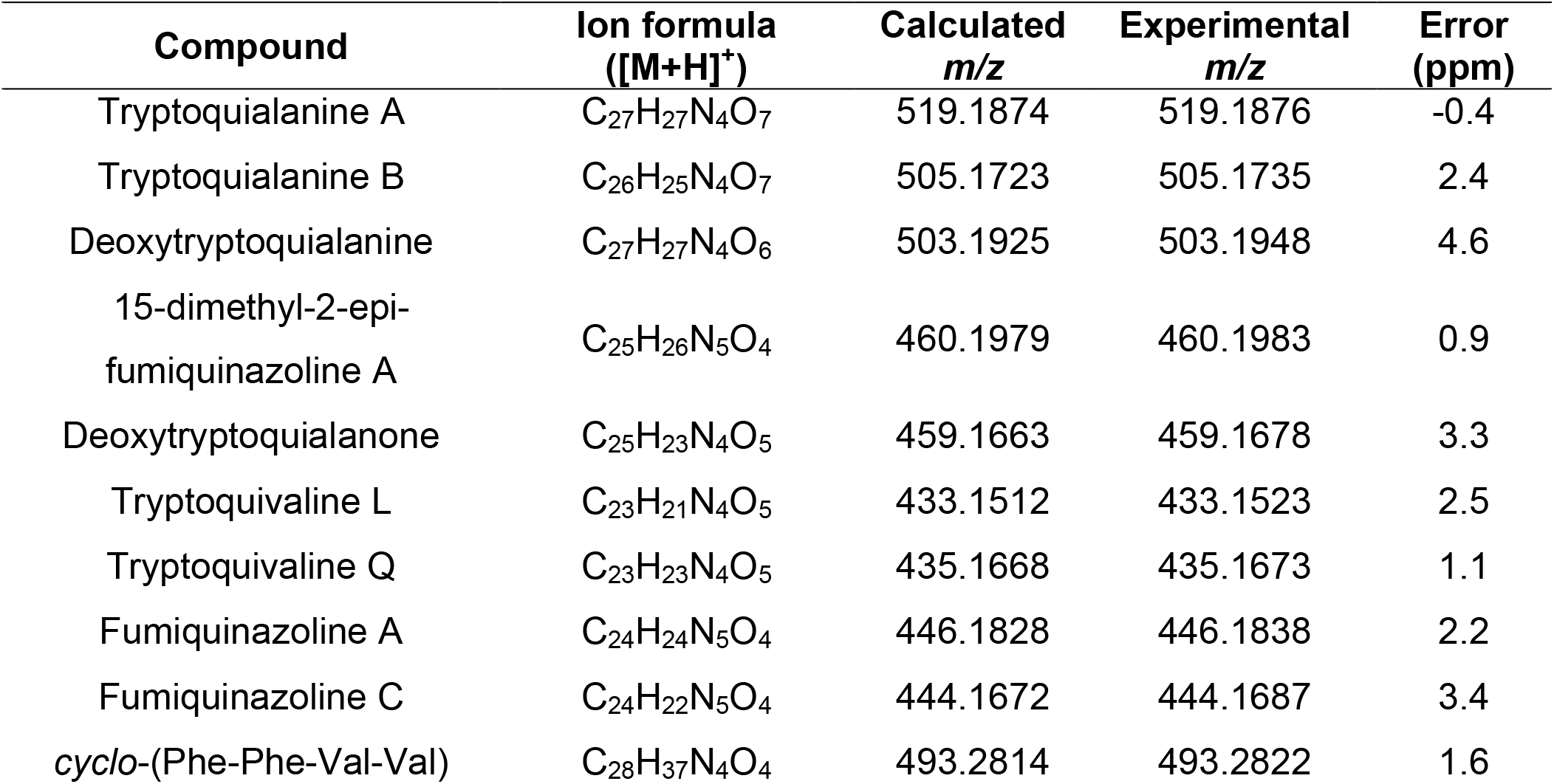
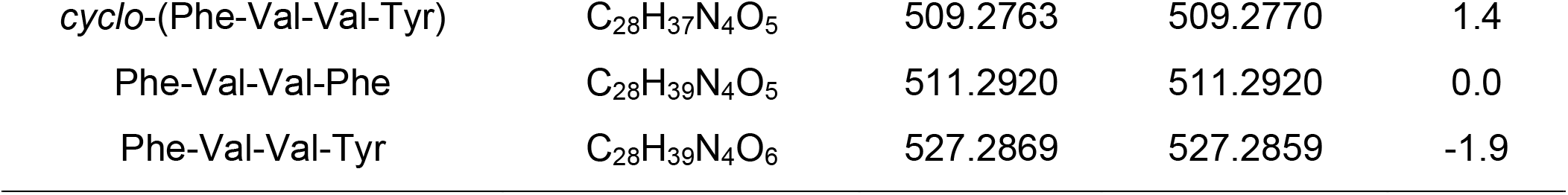
MS data obtained for *P. digitatum* secondary metabolites observed on GNPS molecular network

**FIG 8.**
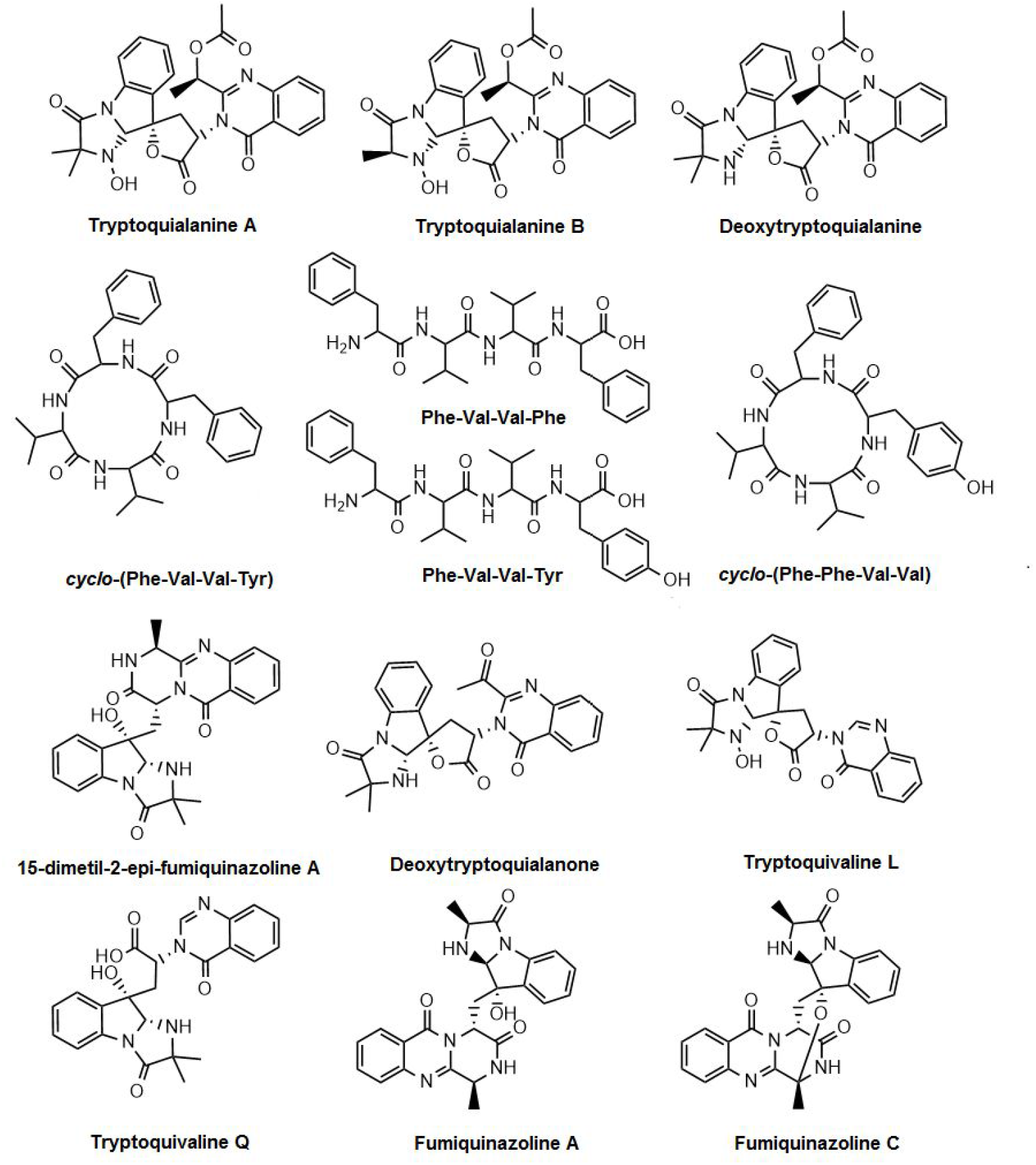
Structures of secondary metabolites identified manually or through the GNPS MS/MS database.

The compounds identified in our analysis included 15-dimethyl-2-*epi*-fumiquinazoline (*m*/*z* 460.19) A, deoxytryptoquialanone (*m*/*z* 459.16), tryptoquivaline L (*m*/*z* 433.15), tryptoquivaline Q (*m*/*z* 435.16 and 417.15), fumiquinazoline A (*m*/*z* 446.18 and 428.16) and fumiquinazoline C (*m*/*z* 444.16 and 426.15). Compounds 15-dimethyl-2-*epi*-fumiquinazoline A and deoxytryptoquialanone are intermediates of the tryptoquialanine biosynthetic pathway (28), while the tryptoquivalines and fumiquinazolines were previously identified as *P. digitatum* metabolites (18). Fungisporin and analogues were also identified in EVs (cluster C, Figure 4). A few differences were observed in clusters D, E, and F (Fig. 7 and 8) considering the production of secondary metabolites by *P. digitatum* in different fruits. All identified compounds were detected in infected plums (at 10 and 13 dpi) and oranges (at 7 dpi) (Fig. S8).

The complete molecular networking obtained for *P. digitatum* is represented in the supporting information (Fig. S9). *P. digitatum* molecular networking was composed by 235 clusters, of which 85 (36 %) were composed of unknown metabolites that were only present in the infected fruits and absent in the control fruits. Molecular networking also showed clusters containing unknown metabolites, present only in infected oranges or only in infected plums (Fig. S10).

## DISCUSSION

Tryptoquialanines are the major secondary metabolites produced by *P. digitatum* (17). The involvement of tryptoquialanines during the infection of citrus fruits by *P. digitatum* was evaluated after deletion of *tqaA* gene (non-ribosomal peptide synthetase) responsible for the biosynthesis of tryptoquialanines. *P. digitatum* mutants deficient in tryptoquialanine A production did not have their virulence affected, when compared to wild type *P. digitatum* cells (15). Thus, tryptoquialanines were initially thought to be dispensable for the pathogenesis in fruits. Other damaging roles could not be ruled out, since they were not investigated in detail. It has been recently reported that TA is accumulated in the citrus surface during the *P. digitatum* pathogenic process (18), suggesting an extracellular export. TA also exhibited insecticidal activity against *Aedes aegypti* larvae (18). These results suggested that tryptoquialanines are involved in the fruit protection against insects that could compete with the fungus for the rotten fruit (18). In co-culture models, it has been observed that *P. digitatum* tryptoquialanines were present in the confrontation zone with citrus pathogens, suggesting that tryptoquialanines participate in antifungal defense mechanisms that could provide competitive advantages during infection of the citrus host (19).

The reports described above and the fact that tryptoquialanines are the major metabolites produced by *P. digitatum* led us to ask if these metabolites would be involved in *P. digitatum* phytotoxic activity. To address this question, we investigated the phytotoxic effects of tryptoquialanines in a seed germination model, as previously established for the evaluation of the phytotoxicity of chemicals (31, 32). This method is simple, sensitive, and of low cost (31). Seed germination is a vulnerable stage in the plant life cycle, during which seedlings are weak, sensitive and more affected by unfavorable conditions (32). Our results indicated that TA was comparable to the herbicide RoundUp® in its ability to inhibit germination. Such inhibitory effect requires the extracellular export of TA, as suggested by its accumulation in the surface of citrus infected with *P. digitatum* (18). We then hypothesized that the transport of the indole alkaloids from *P. digitatum* cells to the extracellular environment would involve EVs, as previously described for fungal proteins, glycans and RNA (26, 33). In our model, EVs were detected in culture and infected citrus fruits. The possibility of co-isolation of plant EVs in the *in vivo* samples cannot be ruled out, since it is well known that plant cells also produce EVs during interaction with fungi (34). However, the similar features of EVs obtained *in vivo* and *in vitro* and our vesicle compositional analysis reinforce the notion that *P. digitatum* produces EVs *in vitro* and during plant infection. The observation of *P. digitatum* EVs gains additional significance considering that most of the studies characterizing fungal EVs used human pathogens as models, which imply that the importance of EV production by phytopathogens has been underscored so far. In this context, it has been only recently demonstrated that EVs from the cotton pathogen *Fusarium oxysporum* f. sp. *vasinfectum* induce a phytotoxic response in plants (26). In the EV cargo of *F. oxysporum* f. sp. v*asinfectum*, 482 enzymes were identified, including two polyketide synthases; yet, the isolated EVs presented a deep purple color, indicating that a naphthoquinone pigment is packaged into the EVs (26). The authors suggested that EVs could be a site of biosynthesis and transport of pigments and other secondary metabolites (26), an idea that is quite complementary to what is presented in our study.

Secondary metabolites participate in the virulence mechanisms of some phytopathogenic fungi, implying that knowledge of metabolite exportation could improve the understanding of the molecular basis of plant infection and fruit protection (4, 16). However, the association of fungal metabolites and EVs has not been established so far in plant infection models. Based on the observation of *P. digitatum* EVs *in vitro* (potato dextrose agar) and *in vivo* (citrus fruits), we identified indole alkaloids and mycotoxins in EV samples. *Penicillium* species are known to produce mycotoxins such as fungisporin (35, 36). Fungisporin and analogues were reported in the cultures of *P. canescens* (29), *P. roqueforti* (30), *P. citrinum* (19) and *P. chrysogenum* (37). Therefore, the production of fungisporin compounds by *P. digitatum* was expected. To the best of our knowledge, the presence of secondary metabolites and mycotoxins in EVs produced by a phytopathogen *in vivo* is herein reported for the first time.

An estimate of the tryptoquialanine levels in EVs could provide insights into the biosynthesis and metabolic flow of these molecules in *P. digitatum*. Previous studies with oranges infected by *P. digitatum* showed that, at 5 days post infection, TA was detected in the orange epicarp, mesocarp and endocarp, with concentrations of 24.810, 388 and 24 µg kg^-1^, respectively (38). TA concentration in the EVs was considerably lower. We then speculate that the biosynthesis of tryptoquialanines may occur in the fungal cells with further export in EVs. This mechanism would differ from that described by Bleackley et al. (2020) in the *F. oxysporum* f. sp. v*asinfectum* EVs (26).

*P. digitatum* EVs induced alterations in the *C. sinensis* seeds, as concluded from the observation of color alteration resulting from tissue lesions. Similar results were reported in recent studies with cotton cotyledons infiltrated with *F. oxysporum* f. sp. V*asinfectum*. In this model, EVs induced discoloration around the sites of infiltration (26). Therefore, the phytotoxic effect observed for the isolated TA was different from that caused by the EVs. These differences were in fact expected, considering that vesicular TA is accompanied by hundreds of other molecules. Those molecules could, for instance, physically interact with TA, altering its relative concentration. In addition, if those additional vesicular molecules have biological effects that differ from that observed for TA alone, it would be very hard to predict what kind of effect would prevail, since the relative concentration of vesicular molecules in the *P. digitatum* model is still unknown. In any case, our results provide a proof-of-concept model showing that *P. digitatum* exports bioactive molecules in EVs that can directly impact the pathogenic process.

*P. digitatum* pathogenesis was believed to be restricted to citrus fruits (13). However, this fungus is also an aggressive pathogen of stone fruits, including nectarines and plums (39, 40). Few studies have investigated the infection of stone fruits by *P. digitatum*. Even though *P. digitatum* disease was characterized at the physical (incidence, lesion diameter, pH) and molecular (gene expression) levels (40, 41), no information on secondary metabolite production has been presented in the literature for this host-pathogen interaction. Since tryptoquialanines and mycotoxins were found in EVs produced during infection, the metabolic profile of *P. digitatum* in different fruits was evaluated, in order to verify if the same metabolites were found in the different hosts. Molecular networking analyses indicate that intermediates of the tryptoquialanine biosynthetic pathway are present in fruits and absent in EVs. This data is in agreement with the quantification level of TA in EVs, reinforcing the idea that tryptoquialanines are only transported by the EVs. Also, our results are the first to identify the production of tryptoquialanines and other indole alkaloids in the *P. digitatum*-stone fruit interaction. Likewise, the similarity between the metabolic profile in the fruits suggests that the production of EVs by *P. digitatum* is not restricted to the citrus fruits, since the same metabolites found in EV cargo obtained from infection in citrus were detected in plums. The clusters containing unknown metabolites present only in infected oranges or only in infected plums (Fig. S9) suggest that the metabolite production of *P. digitatum* can vary depending on the infected fruit.

## CONCLUSIONS

This work is the first to report that *P. digitatum* is able to release EVs, and to report secondary metabolites in EVs produced by a phytopathogen *in vivo*. Furthermore, we suggested that TA is synthesized intracellularly and exported in EVs. Molecular networking confirmed our hypothesis that tryptoquialanines and mycotoxins are delivered through EVs during the infection process, since the intermediates of the tryptoquialanine biosynthetic pathway are absent in the EVs. This delivery system is not restricted to citrus, and occurs in different types of fruits, such as plums.

A novel phytotoxic function for *P. digitatum* EVs and for tryptoquialanines was observed. EVs caused alterations in the physiology of *C. sinensis* seeds tissues, while TA inhibited 100 % of seed germination. The presence of alkaloids and mycotoxins in phytotoxic EVs opens new venues for the investigation of fungal secretion and its relationship with plant pathogenesis. Also, our results provided new insights into the biological role of the indole alkaloids and the infection strategies used by the phytopathogen *P. digitatum*.

## MATERIALS AND METHODS

### Fungal strain and culture conditions

*P. digitatum* strain is deposited in the Spanish Type Culture Collection (CECT) (accession code: CECT20796). The fungus was cultured in commercial potato dextrose agar (PDA) (darkness, 7 days at 25 °C). Conidial suspensions were prepared in sterile distilled water and adjusted to a final concentration of 1.0 × 10^6^ conidia mL^-1^.

### Purification of tryptoquialanine A by HPLC

*P. digitatum* was cultivated in 12 L of PDA distributed in Petri dishes. After cultivation, the content of the Petri dishes was sliced and transferred to Erlenmeyer flasks. The content of Erlenmeyer flasks was extracted twice with ethyl acetate (EtOAc) under sonication in an ultrasonic bath during 1 h. The mixture of agar, mycelia, and EtOAc was filtered and the solvent was removed under reduced pressure.

The *P. digitatum* EtOAc extract was suspended in methanol (MeOH), filtered, and subjected to a separation by high-performance liquid chromatography (HPLC) in order to obtain pure tryptoquialanine A. HPLC separation was performed with a Phenomenex column Luna 5 µm Phenyl-Hexyl (250 × 4.6 mm) using a SHIMADZU prominence HPLC LC-20AT instrument connected to a CBM-20A Communication Bus Module, to a SPD-M20A photodiode array detector and to a SIL-20A auto sampler. The mobile phases were 0.1% (v/v) formic acid in water (A) and acetonitrile (B). Flow rate was 1.0 mL min^-1^. Elution was performed as follows (A:B): gradient from 95:5 up to 55:45 during 30 min, then up to 35:65 from 30 to 52 min, then up to 5:95 from 52 to 55 min remaining in this condition for 5 min. Column reconditioning between each injection was a gradient to 95:5 from 60 to 61 min remaining in this condition for 9 min. Semi preparative HPLC separations were performed with a Phenomenex column Luna 5 µm Phenyl-Hexyl (250 × 10 mm) using a Waters 1525 Binary HPLC Pump equipped with Waters 2998 Photodiode Array Detector and Waters Fraction Collector III. The eluent was the same as indicated above with a flow rate of 4.7 mL min^-1^

### Seed germination test (phytotoxicity assay)

The phytotoxicity of tryptoquialanine A on seed germination was evaluated as previously described with a few modifications (42–45). Briefly, *C. sinensis* seeds were manually collected from oranges purchased at a local grocery store (Campinas, SP, Brazil). Seeds coats were removed, and seeds were immersed in a 50 % (v/v) commercial bleach solution during 15 min for surface sterilization. Six sterilized seeds were placed in each Petri dish (6 cm) lined with two filter papers. A volume of 2.5 mL of treatment solutions were added to the plate. As negative control (NC), seeds were treated with sterile distilled H_2_O containing dimethyl sulfoxide (DMSO) 3 % (v/v). Tryptoquialanine A (TA) was solubilized in DMSO and diluted in sterile distilled water to a final concentration of 500, 1.000, and 3.000 ppm. The commercial herbicide RoundUP® was utilized as a positive control (PC), diluted to the concentrations of 10.000 and 3.000 ppm in sterile distilled water containing DMSO 3% (v/v). Treatment solutions were filtered through 0.22 µm membranes. Petri dishes were sealed with tape and incubated in a biochemical oxygen demand (BOD) chamber at 25 °C with photoperiods of 12 h during 10 days. After incubation, the percentage of seed germination was calculated as described in Equation 1, considering complete, proportionate, and healthy development.

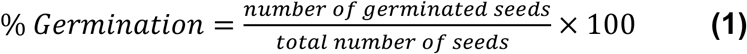

To evaluate the phytotoxic activity of EVs, uncoated and sterilized *C. sinensis* seeds were placed in a 24-well cell culture plate lined with filter papers (1 seed per well). The seeds were treated with 100 µl of a PBS solution of *P. digitatum* EVs (2.1 × 10^10^ EVs mL^-1^). Negative controls (NC) were performed using 100 µl of PBS and, for positive controls (PC), 100 µl of herbicide RoundUP® diluted in PBS (10.000 ppm) was used. The plate was sealed and incubated as described above.

### Infection of fruits by *P. digitatum* (*in vivo* assays) and metabolite extraction

For *in vivo* assays, mature oranges (*C. sinensis*) and plums (*Prunus salicina*) obtained from a local grocery store (Campinas, SP, Brazil) were surfaced sterilized and wounded (18). Four fruits (2 oranges and 2 plums) were infected with 15 µL of a *P. digitatum* 1.0 × 10^6^ conidia mL^-1^ solution. Control fruits (2 oranges and 2 plums) were also included. Infected and control fruits were stored in sterile 500 mL beakers, in darkness at 25 °C. The fruits were incubated for different days post inoculation (dpi) in triplicates.

After the infection period (7 dpi for oranges, 10 and 13 dpi for plums), extraction of infected fruits was performed as previously described, with few modifications (46). Fruits were cut around the infected area (4 cm x 4 cm) and collected fruit pieces were extracted with 5 mL of MeOH during 1 h in ultrasonic bath. Same procedure was performed for control fruits. MeOH extracts were filtered, dried with a N_2_ flux, and stored at −20 °C.

### Isolation of *P. digitatum* EVs and metabolite extraction

Isolation of *P. digitatum* EVs produced *in vitro* was performed as previously described, with a few modifications (27). Fungal cells were cultivated and softly scrapped from PDA plates (triplicates, 20 mL of PDA per plate) using a sterile spatula. Fungal mycelia were transferred to a Falcon tube filled up with 30 mL of sterile Phosphate-Buffered Saline (PBS). For the analysis of EVs *in vivo*, nine oranges (*C. sinensis*) were infected with *P. digitatum* (section 2.4). Infected fruits were incubated for 7 days (darkness, 25 °C). Then, fungal cells in the infected areas of fruits were softly scrapped using a sterile spatula and transferred to a Falcon tube filled up with 30 mL of PBS. 30 mL cell suspensions obtained *in vivo* or *in vitro* were sequentially centrifuged to remove fungal cells (5,000 x *g* for 15 min at 4 °C) and possible debris (15,000 x *g* for 15 min at 4 °C). The remaining supernatants were filtered through 0.45-μm-pore syringe filters and ultracentrifuged to collect EVs (100,000 x *g* for 1 h at 4 °C). Ultracentrifugation pellets were negatively stained and analyzed by transmission electron microscopy (TEM) as previously described (27). Briefly, EVs samples were transferred to carbon- and Formvar-coated grids and negatively stained with 1 % (v/v) uranyl acetate for 10 min. The grids were then blotted dry before immediately being observed in a JEOL 1400Plus transmission electron microscope at 90 kV. The same samples were submitted to nanoparticle tracking analysis (NTA) on an LM10 nanoparticle analysis system, coupled with a 488-nm laser and equipped with an _S_CMOS camera and a syringe pump (Malvern Panalytical, Malvern, United Kingdom). Recorded data were acquired and analyzed using the NTA 3.0 software (Malvern Panalytical).

To study the vesicular cargo, EVs obtained *in vivo* were extracted with 1 mL of MeOH HPLC grade during 1 h in an ultrasonic bath.

### Mass spectrometry (MS) analyses

#### *In vivo* extracts

*In vivo* extracts were resuspended in 1 mL of MeOH HPLC grade. An aliquot of 100 μL was diluted in 900 µL of MeOH HPLC grade, filtered through 0.22 µm membranes, and collected in glass vials. UHPLC-MS analyses were performed in a Waters Acquity UPLC H-Class chromatograph coupled to a Waters Xevo G2-XS QToF mass spectrometer using electrospray ionization. Conditions: positive mode, capillary voltage at 1.2 kV; source temperature at 100 °C; cone gas (N_2_) flow of 50 L h-1; desolvation gas (N_2_) flow of 750 L h^-1^ and *m*/*z* range of 100-1500. MS/MS analyses were performed using collision energy ramp of 6-9 V (low mass) and 60-80 V (high mass). A BEH C18 column (2.1 mm x 100 mm x 1.7 µm) was used. Mobile phases were 0.1 % (v/v) formic acid in water (A) and acetonitrile (B). Eluent profile (A:B) 0-6 min, gradient from 90:10 up to 50:50; 6-9 min, gradient up to 2:98; 9-10 min, gradient up to 90:10. Flow rate was 0.2 mL min^-1^. Injection volume was 2 µL. Operation and spectra analysis were conducted using Waters MassLynx V.4.1. software.

#### *P. digitatum* EV extracts

1 mL of EV extracts was filtered through 0.22 µm membranes into glass vials. UHPLC-MS analysis were performed using a Thermo Scientific QExactive® hybrid Quadrupole-Orbitrap mass spectrometer with the following parameters: electrospray ionization in positive mode, capillary voltage at +3.5 kV; capillary temperature at 250 °C; S-lens of 50 V and *m*/*z* range of 133.40-2000.00. MS/MS were performed using normalized collision energy (NCE) of 30 eV and 5 precursors per cycle were selected. Stationary phase: Thermo Scientific Accucore C18 2.6 µm (2.1 mm x 100 mm) column. Mobile phases were 0.1 % (v/v) formic acid in water (A) and acetonitrile (B). Eluent profile (A:B) 0-10 min, gradient from 95:5 up to 2:98; held for 5 min; 15-16.2 min gradient up to 95:5; held for 8.8 min. Flow rate was 0.2 mL min^-1^. Injection volume was 3 µL. Operation and spectra analysis were conducted using Xcalibur software (version 3.0.63) developed by Thermo Fisher Scientific.

### Seed extracts

Two seeds of each condition, TA (3.000 ppm), PC (10.000 ppm) and NC, were macerated with liquid nitrogen, in triplicate. Aliquots of 100 mg of macerated seeds were extracted in plastic tubes with 2 mL of MeOH containing 0.1 % (v/v) formic acid during 1 h in an ultrasonic bath. The extracts were filtered (0.22 µm), dried with a N_2_ flux, and stored at −20 °C.

Seed extracts were resuspended in 1 mL of MeOH and aliquots of 100 µL and diluted with 900 µL and filtered through a 0.22 µm membrane. UHPLC-MS analyses were performed using a Thermo Scientific QExactive® hybrid Quadrupole-Orbitrap mass spectrometer with the following parameters: electrospray ionization in positive mode, capillary voltage at 3.5 kV; capillary temperature at 300 °C; S-lens of 50 V and *m*/*z* range of 100.00-1500.00. MS/MS were performed using normalized collision energy (NCE) of 20, 30 and 40 eV and maximum 5 precursors per cycle were selected. A Waters ACQUITY UPLC® BEH C18 1.7 µm (2.1 mm x 50 mm) column was used. Mobile phases were 0.1 % (v/v) formic acid in water (A) and acetonitrile (B). Eluent profile (A:B) 0-10 min, gradient from 95:5 up to 2:98; held for 5 min; 15-16.2 min gradient up to 95:5; held for 3.8 min. Flow rate was 0.2 mL min^-1^. UHPLC-MS operation and spectra analyses were performed using Xcalibur software (version 3.0.63). Samples were injected in random order. A Quality Control (QC) sample was prepared with 50 µL of each sample and was injected three times at the beginning of the batch and after three sample injections (47–49).

### Quantification of tryptoquialanine A

Standard TA isolated from *P. digitatum* and EVs extract were analyzed using a Waters ACQUITY UPLC system coupled to Waters Micromass Quattro Micro TM API, with electrospray ionization source and *triple quadrupole* mass *analyzer*. Analyses were performed in the positive mode with *m*/*z* range of 100-1200, capillary voltage of 3 kV, cone voltage of 25 V, inlet capillary temperature of 150 °C and nebulizing gas temperature of 200 °C. Stationary phase: Thermo Scientific column Accucore C18 2.6 µm (2.1 mm x 100 mm). Mobile phase: 0.1% formic acid (A) and acetonitrile (B). Eluent profile (A/B): 95/5 up to 2/98 within 10 min, hold for 5 min, up to 95/5 within 1.2 min and hold for 3.8 min. The total run time was 20 min for each run and the flow rate was 0.2 mL min^-1^. Injection volume: 10 µL. All the operation and spectra analysis were conducted using Waters MassLynx V.4.1.

For the construction of the calibration curve, standard TA was diluted in the range concentration of 6.25 to 0.006 µg mL^-1^, and selected reaction monitoring (SRM) analyses were performed following conditions as previously described: *m*/*z* 519 → 197 (quantification) and *m*/*z* 519 → 213 (monitoring), collision energy of 22 eV (38).

For quantification of tryptoquialanine A in EVs, 40 µL of a 2.1 × 10^10^ EVs mL^-1^ solution was dried and extracted with 100 µL of MeOH HPLC grade as previously described. 100 µL of EVs extract solution was transferred to glass vial and analyses were performed in duplicate.

### Statistical and metabolomic analysis

Feature detection was performed on XCMS online (version 3.5.1) using the following parameters: method: centWave, prefilter peaks and intensity: 3 and 5000, ppm: 2.5, Signal/Noise threshold: 10, peak width: 5-20, mzdiff: 0.01 and noise filter: 1000. Preprocessing included median fold change normalization on XCMS Online. Multivariate and univariate analyses of the feature list were performed with the MetaboAnalyst tool (version 4.0). Pareto scaling was applied. One-way analysis of variance (ANOVA) was performed and all the results were analyzed using a confidence level of 95 % and significance level corresponding to *p* < 0.05. Principal component analysis (PCA) was performed for an exploratory analysis, followed by partial least squares discriminant analysis (PLS-DA). A permutation test (cross validation) was performed to determine the reliability of the created PLS-DA model.

### Molecular networking analyses

MS data were converted to mzXML format using MSConvert GUI, a tool of ProteoWizard package. Molecular networks for *in vivo* assays and EVs extracts were created using the mzXML files on the online workflow at the Global Natural Products Social Molecular Networking (GNPS) platform (http://gnps.ucsd.edu). Data was filtered by removing all MS/MS peaks within +/- 17 Da of the precursor ion. MS/MS spectra were window filtered by choosing only the top 6 peaks in the +/- 50 Da window throughout the spectrum. The data was then clustered with MS-Cluster with a parent mass tolerance of 0.02 Da and a MS/MS fragment ion tolerance of 0.02 Da to create consensus spectra. Consensus spectra that contained less than 2 spectra were discarded. A network was then created where edges were filtered to have a cosine score above 0.6 and more than 5 matched peaks. The spectra in the network were then searched against GNPS’ spectral libraries. The library spectra were filtered in the same manner as the input data. All matches between network spectra and library spectra were required to have a score above 0.6 and at least 5 matched peaks (50).

## ACKNOWLEDGMENTS

This study was financed in part by the Coordenação de Aperfeiçoamento de Pessoal de Nível Superior - Brasil (CAPES) - Finance Code 001, Fundação de Amparo a Pesquisa no Estado de São Paulo [grants number 2019/11563-2, 2019/06359-7, 2017/24462-4, 2013/50228-8 and 2015/01017-0] L’Oréal Brazil, together with ABC and UNESCO in Brazil. M.L.R. was supported by grants from the Brazilian Ministry of Health (grant 440015/ 2018-9), Conselho Nacional de Desenvolvimento Científico e Tecnológico (CNPq; grants 405520/2018-2 and 301304/2017-3), and Fiocruz (grants PROEP-ICC 442186/2019-3, VPPCB-007-FIO-18, and VPPIS-001-FIO18). We also acknowledge support from the Instituto Nacional de Ciência e Tecnologia de Inovação em Doenças de Populações Negligenciadas (INCT-IDPN).

## SUPPLEMENTAL MATERIAL

**FIG S1** HPLC chromatograms (280 nm) for (A) *P. digitatum* extract and (B) purified tryptoquialanine A. HPLC-MS analysis of purified tryptoquialanine A (*m*/*z* 519): (C) extracted ion chromatogram and (D) mass spectrum.

**FIG S2** Extracted ion chromatograms of *m*/*z* 519.18 for (A) *P. digitatum* EVs extract, (B) MS analysis control and (C) EV isolation methodology control. (D) Mass spectrum of ion [M+H]^+^ *m*/*z* 519.1877 obtained for tryptoquialanine A (error = 0.61 ppm) at 9.2 min. Extracted ion chromatograms of *m*/*z* 505.17 for (E) *P. digitatum* EV extract, (F) MS analysis control and (G) EV isolation methodology control. (H) Mass spectrum of ion [M+H]^+^ *m*/*z* 505.1719 obtained for tryptoquialanine B (error = 0.26 ppm) at 8.7 min. Extracted ion chromatograms of *m*/*z* 503.19 for (I) *P. digitatum* EV extract, (J) MS analysis control and (K) EV isolation methodology control. (L) Mass spectrum of ion [M+H]^+^ *m*/*z* 505.1926 obtained for deoxytryptoquialanine (error = 0.23 ppm) at 8.6 min.

**FIG S3** Extracted ion chromatograms of *m*/*z* 509.27 for (A) *P. digitatum* EV extract, (B) MS analysis control and (C) EV isolation methodology control. (D) Mass spectrum of ion [M+H]^+^ *m*/*z* 509.2761 obtained for *cyclo*-(Phe-Val-Val-Tyr) (error = 0.48 ppm) at 8.6 min. Extracted ion chromatograms of *m*/*z* 511.29 for (E) *P. digitatum* EV extract, (F) MS analysis control and (G) EV isolation methodology control. (H) Mass spectrum of ion [M+H]^+^ *m*/*z* 511.2917 obtained for Phe-Val-Val-Phe (error = 0.43 ppm) at 7.1 min. Extracted ion chromatograms of *m*/*z* 527.28 for (I) *P. digitatum* EV extract, (J) MS analysis control and (K) EV isolation methodology control. (L) Mass spectrum of ion [M+H]^+^ *m*/*z* 527.2866 obtained for Phe-Val-Val-Tyr (error = 0.40 ppm) at 6.4 min. Extracted ion chromatograms of *m*/*z* 493.28 for (M) *P. digitatum* EV extract, (N) MS analysis control and (O) EV isolation methodology control. (P) Mass spectrum of ion [M+H]^+^ *m*/*z* 493.2809 obtained for *cyclo*-(Phe-Phe-Val-Val) (error = 0.03 ppm) at 9.7 min.

**FIG S4** Comparison of MS/MS spectra between (A) *P. digitatum* EV extract and (B) isolated tryptoquialanine A, (C) *P. digitatum* EV extract and (D) isolated tryptoquialanine B, (E) *P. digitatum* EV extract and (F) isolated deoxytryptoquialanine.

**FIG S5** (A) MS/MS match between GNPS database (green) and compound *cyclo*-(Phe-Phe-Val-Val) (black). MS/MS spectrum of (B) *m*/*z* 511.29 and (C) *m*/*z* 527.28 obtained from *P. digitatum* EV extract. Fragmentation pattern was compared with MS/MS data of compounds Phe-Val-Val-Phe and Phe-Val-Val-Tyr reported in the literature.

**FIG S6** UHPLC-MS/MS calibration curve obtained for tryptoquialanine A.

**FIG S7** Mass spectrum and MS/MS spectrum for (A) tryptoquialanine A ion [M+H]^+^ *m*/*z* 519.1876, (B) tryptoquialanine B ion [M+H]^+^ *m*/*z* 505.1735, (C) deoxytryptoquialanine ion [M+H]^+^ *m*/*z* 503.1948, (D) *cyclo*-(Phe-Val-Val-Tyr) ion [M+H]^+^ *m*/*z* 509.2770, (E) Phe-Val-Val-Phe ion [M+H]^+^ *m*/*z* 511.2920, (F) Phe-Val-Val-Tyr ion [M+H]^+^ *m*/*z* 527.2859, (G) *cyclo*-(Phe-Phe-Val-Val) ion [M+H]^+^ *m*/*z* 493.2822, (H) 15-dimethyl-2-epi-fumiquinazoline A ion [M+H]^+^ *m*/*z* 460.1983, (I) deoxytryptoquialanone ion [M+H]^+^ *m*/*z* 459.1678, (J) tryptoquivaline L ion [M+H]^+^ *m*/*z* 433.1523, (K) tryptoquivaline Q ion [M+H]^+^ *m*/*z* 435.1673, (L) fumiquinazoline A ion [M+H]^+^ *m*/*z* 446.1838 and (M) fumiquinazoline C ion [M+H]^+^ *m*/*z* 444.1687.

**FIG S8** Fruits infected with *P. digitatum*. (A) plum at 10 dpi, (B) plum at 13 dpi and (C) orange at 7 dpi.

**FIG S9** Complete molecular networking obtained for extracts of plums and citrus fruits infected by *P. digitatum*.

**FIG S10** Examples of clusters obtained by molecular networking for extracts of *P. digitatum* infection in different fruits. The unknown compounds are present just in one type of the fruits and absent in the control fruits

